# A new *Betaflexivirus* species in *Ferraria crispa* plants

**DOI:** 10.1101/2024.09.02.610804

**Authors:** Rob J. Dekker, Wim C. de Leeuw, Marina van Olst, Wim A. Ensink, Selina van Leeuwen, Timo M. Breit, Martijs J. Jonker

## Abstract

The global virome is still largely unknown. In this study we describe the discovery of a new *Betaflexivirus* species in *Ferraria crispa* plants. The plant samples were collected in an experiment of 25 *Asparagales* plants, obtained from an urban botanic garden in the Netherlands, and analyzed by smallRNA-seq as well as RNA-seq. The new *Betaflexivirus* only occurred in four *Ferraria cripsa* plants and was tentatively named Ferraria Betaflexivirus 1 (FerBfV-1). This *Betaflexivirus* showed an RNA genome structure characteristic for the genus *Capillovirus*, a single-stranded (+) RNA virus. The closest known *Capilloviruses* in GenBank showed a protein similarity less than 49%, which is well below the proposed demarcation value of species in this genus. Further comparison analysis of the *Capillovirus* genus revealed the possible presence of two groups.

## Introduction

As far as we know, all living organisms must battle infections by one or more virus species (Greene and Reid, 2013). Hence, viruses play a crucial role in life on earth. However, aside from human pathogenic viruses, other viruses, including phytoviruses, are underrepresented in our life sciences studies and knowledge. This may be a consequence of the paradigm that viruses are not considered living organisms and thus have no place in “the tree of life” (Moreira and López-García, 2009). Yet, since viruses significantly impact the lives of living organisms, it is imperative to understand their functioning. For this, it is important to identify as many viruses as possible from the world-wide virome (Ignacio-Espinoza *et al*., 2013). Fortunately, advancements in high-throughput sequencing technologies have greatly facilitated the discovery of viruses in all sorts of organisms (Villamor *et al*., 2019).

Our research focus includes phytoviruses, and in our pursuit for new viruses, we wanted to investigate if urban botanic gardens, with their diverse collection of exotic plants, may serve as reservoirs for phytoviruses. To enhance our understanding of the global plant virome’s complexity, we conducted a study on 25 *Asparagales* plants, sourced directly from a botanical garden in Amsterdam, The Netherlands. Since most phytoviruses invoke an siRNA response in an infected plant, we sequenced the RNA of the plant samples using both smallRNA-seq and RNA-seq techniques.

In this report we describe a new single-stranded (+) RNA *Betaflexivirus* from the genus *Capillovirus* (Murphy *et al. 1995*, Magome *et al*. 1997) that was present in several *Ferrari crispa* plants and is distinctly different from several recently reported *Betaflexiviruses* (Marais *et al*., 2018; Roberts *et al*., 2018; Li *et al*., 2020; He *et al*., 2023; Khan *et al*., 2024). This new virus was identified alongside two other newly discovered viruses in the same set of samples (Dekker et al., 2024a and 2024b).

## Material and methods

### Samples

Samples of leaves from 25 *Asparagales* plants were collected from Hortus Botanicus, a botanic garden in Amsterdam, the Netherlands, on February 14, 2019. Details about the plant genera can be found in Supplemental Table ST1.

### RNA isolation

Small-RNA was isolated by grinding a flash-frozen ±1 cm^2^ leaf fragment to fine powder using mortar and pestle, dissolving the powder in QIAzol Lysis Reagent (Qiagen) and purifying the RNA using the miRNeasy Mini Kit (Qiagen). Separation of the total RNA in a small (<200 nt) and large (>200 nt) fraction, including DNase treatment of the large RNA isolates, was performed as described in the manufacturer’s instructions. The concentration of the RNA was determined using a NanoDrop ND-2000 (Thermo Fisher Scientific) and RNA quality was assessed using the 2200 TapeStation System with Agilent RNA ScreenTapes (Agilent Technologies).

### RNA-sequencing

Barcoded smallRNA-seq and RNA-seq libraries were generated using a Small RNA-seq Library Prep Kit (Lexogen) and a TruSeq Stranded Total RNA with Ribo-Zero Plant kit (Illumina), respectively. The size distribution of the libraries with indexed adapters was assessed using a 2200 TapeStation System with Agilent D1000 ScreenTapes (Agilent Technologies). The smallRNA-seq libraries from samples S01 to S12 and from samples S14-S26 were clustered and sequenced at 2×75 bp and 1×75 bp, on a NextSeq 550 System using a NextSeq 500/550 Mid Output Kit v2.5 or a NextSeq 500/550 High Output Kit v2.5 (75 cycles or 150 cycles; Illumina), respectively. RNA-seq libraries were clustered and sequenced at 2×150 bp on a NovaSeq 6000 System using the NovaSeq 6000 S4 Reagent Kit v1.5 (300 cycles; Illumina).

### Bioinformatics analyses

Sequencing reads were trimmed using trimmomatic v0.39 (Bolger *et al*., 2014) [parameters: LEADING:3; TRAILING:3; SLIDINGWINDOW:4:15; MINLEN:19]. Mapping of the trimmed reads to the NCBI virus database was performed using Bowtie2 v2.4.1 (Langmead *et al*., 2012). Contigs were assembled from smallRNA-seq data using all trimmed reads as input for SPAdes De Novo Assembler (Prjibelski *et al*., 2020) with parameter settings: only-assembler mode, coverage depth cutoff 10, and kmer length 17, 18 and 21. Assembly of contigs from RNA-seq data was performed with default settings.

Scanning of contig sequences for potential RdRp-like proteins was performed using PalmScan (Babaian *et al*., 2022) and LucaProt (Hou *et al*., 2023).

## Results and discussion

In our pursuit to discover yet unknown plant viruses, we analyzed 25 *Asparagales* plants covering six different genera from an urban botanic garden by smallRNA-seq and RNA-seq. For both deep-sequencing techniques, sufficient reads were obtained to start the bioinformatics virus-discovery exploration. In sample S16, analysis of both sequencing datasets revealed several RNA sequence contigs suggesting the presence of a potential new Betaflexivirus. After the final sequence assembly, we discovered a continuous RNA sequence of 6,313 nucleotides with a poly-A tail, displaying all the characteristic features of the *Betaflexiviridae* family (Figure 1; Adams *et al*., 2012).

**Figure 1.**
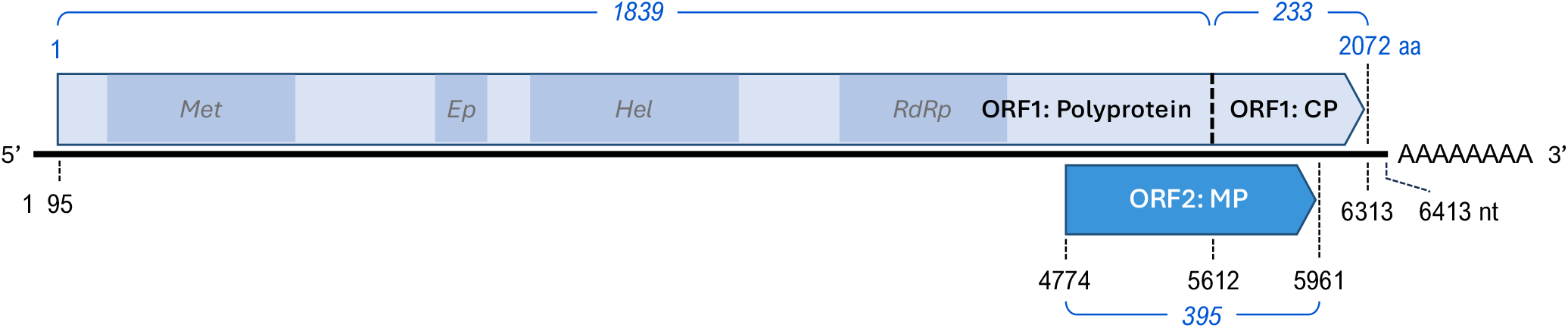
Schematic representation of the genome and proteins of the Ferraria betaflexivirus 1 virus. Indicated are the positions of the proposed polyprotein cleavage site for the coat protein (CP) and the translation start and stop sites for ORF1 (Polyprotein-CP) and ORF2 (movement protein, MP) of the Ferraria betaflexivirus 1 (FerBfV-1). Sizes of the mature protein are indicated in blue italics. Four putative conserved protein domains in ORF1 are shown in darker blue: viral methyltransferase (pfam 01660, *Met*); Carlavirus endopeptidase (pfam PF05379, *Ep*); Viral (Superfamily 1) RNA helicase (pfam PF01443, *Hel*); and RNA dependent RNA polymerase (pfam PF00978, *RdRp*). The complete genomic RNA sequence is deposited in NCBI GenBank (PQ073131).

There appear to be two overlapping open reading frames (ORFs) oriented in the same direction: ORF1 (2,071 amino acids) encodes a polyprotein-coat protein, and ORF2 (395 amino acids) encodes a putative movement protein (Figure 1). Several conserved protein domains, such as RNA dependent RNA polymerase (RdRp) were recognized in ORF1: Polyprotein.

Comparison of the new *Betaflexivirus* protein sequences with all GenBank protein sequences revealed weak similarity to known *Capilloviruses*, a genus within the *Betaflexiviridae* family. The most similar known virus is Rhodiola Betaflexivirus 1, showing 48.5% similarity at the protein level and 67.3% similarity at the RNA level, but with only 33% coverage (Table 1). Both similarity values fall well below the proposed species demarcation thresholds for the *Capillovirus* genus, which require less than approximately 72% nucleotide identity or 80% amino acid identity between their coat protein or polymerase genes. (Adams *et al*., 2012; Silva *et al*., 2022). Therefore, this new *Capillovirus* represents a new species within the *Betaflexivirus* genus.

**Table 1.**
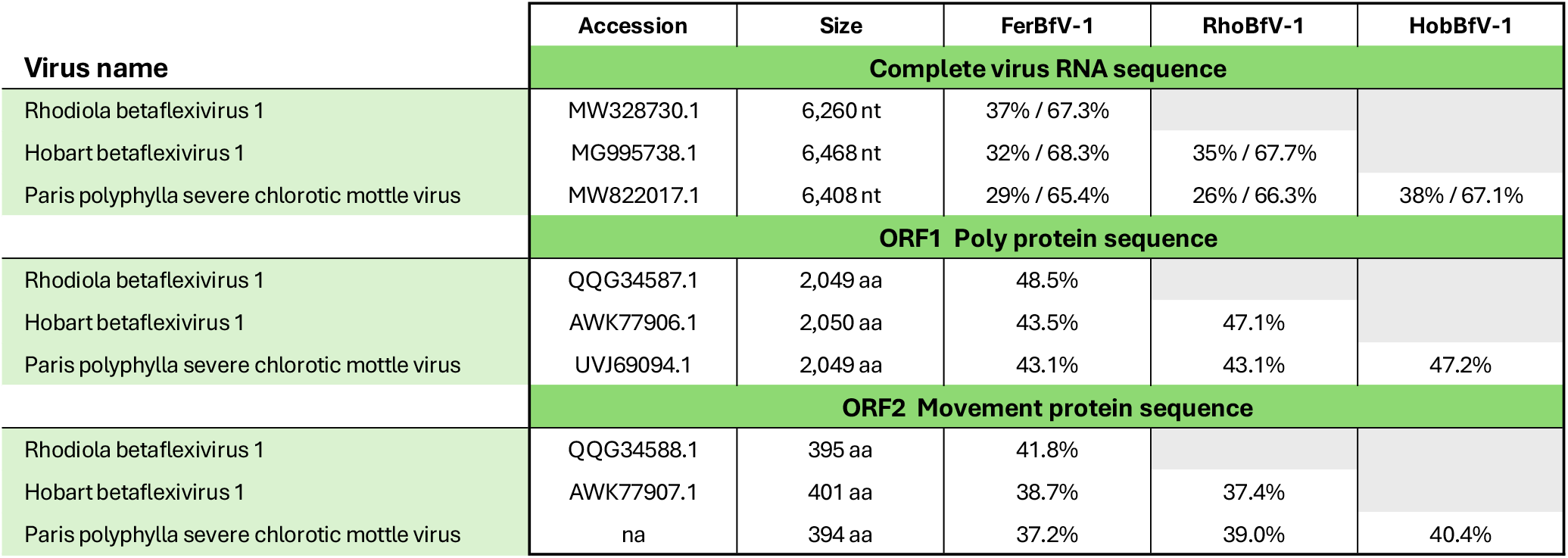
Comparative analysis of RNA and protein sequences of FerBfV-1 and three closely related viruses. Comparison of the RNA and protein sequences of FerBfV-1 with three closely related viruses: Rhodiola betaflexivirus 1 (RhoBfV-1), Hobart betaflexivirus 1 (HobBfV-1), and Paris polyphylla severe chlorotic mottle virus (PpSCMV), obtained using the indicated NCBI GenBank accession numbers. The similarity of the complete viral genomes is expressed as ORF1/ORF2 (%).

Backmapping the smallRNA-seq and RNA-seq reads of all samples to the new *Betaflexivirus*, clearly showed that this virus was only present in four samples (Supplemental Table ST2). Since all these samples were collected from *Ferraria crispa* plants, we have tentatively named this new virus Ferraria betaflexivirus 1 (FerBfV-1).

Building on the *Capillovirus* comparison by He *et al*. (2023), we constructed a phylogenetic tree using the complete ORF1 protein sequences from representative members of the *Capillovirus* species, other *Betaflexivirus* genera, and FerBfV-1 (Figure 2). As expected, FerBfV-1 clustered with other *Capilloviruses*. Additionally, we observed that the *Capillovirus* genus appears to divide into two distinct groups (Figure 2, Supplemental Table ST2). This was supported by protein sequence alignments of the N-termini and the RdRp motifs A, B, and C (Jia and Gong, 2019) from representative members of all *Capillovirus* species (Figure 3), which revealed distinct, conserved differences between the two *Capillovirus* groups.

**Figure 2.**
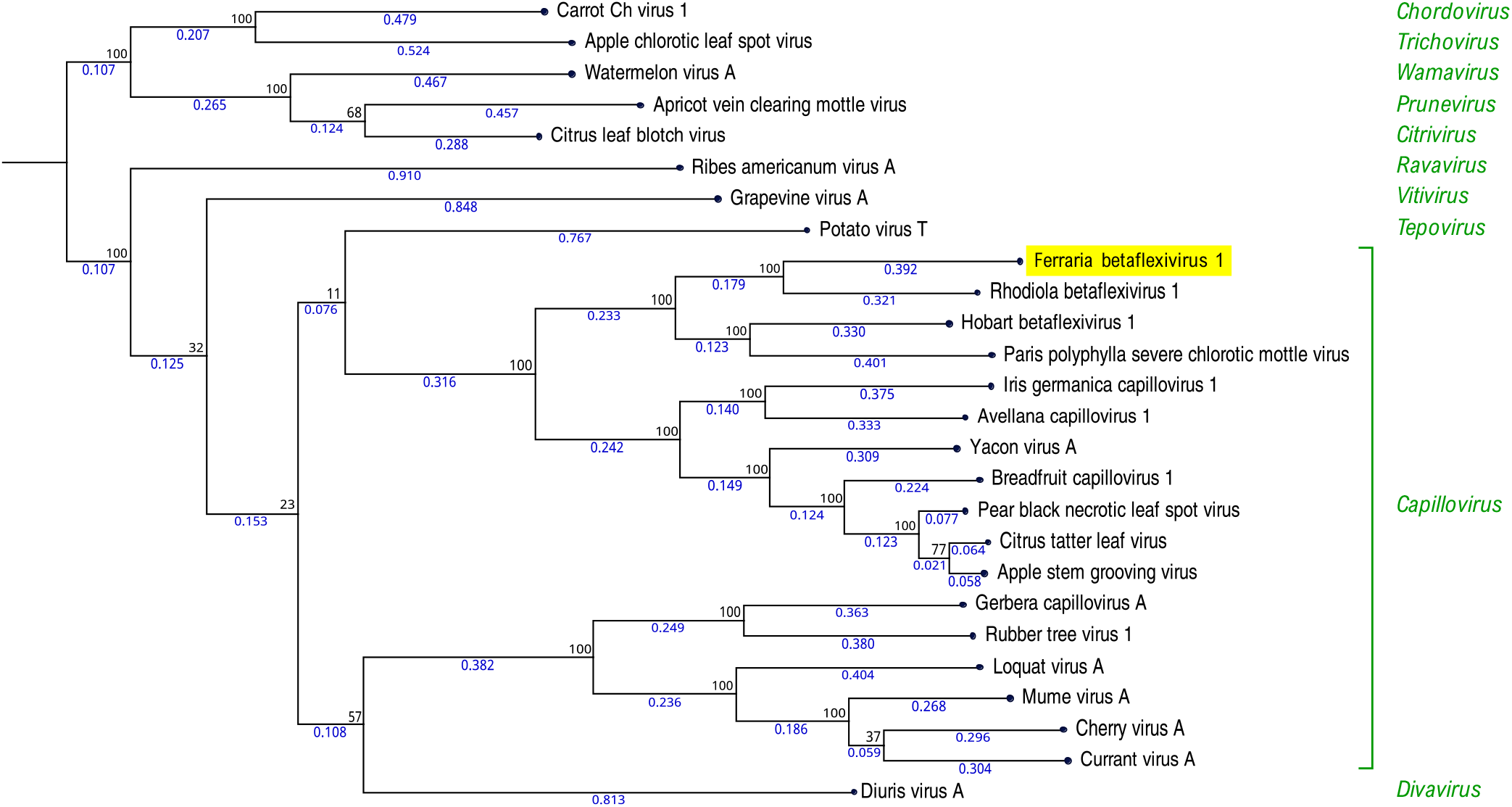
Phylogenetic analysis of Ferraria Betaflexivirus 1 ORF1 protein. Ferraria betaflexivirus 1 (FerBfV-1), identified in this study (highlighted in yellow), is shown alongside 25 species from 10 different genera within the family *Betaflexiviridae* (highlighted in green) sourced from GenBank (see accession numbers in Supplementary Table ST2). The tree was constructed using CLC Genomics Workbench v24.0.1. Initially, amino acid sequences of the ORF1 protein were aligned with gap cost settings: open 5, extension 1, and end gap ‘Free’. The CLC’s Maximum Likelihood Phylogeny 1.4 tool was then used to construct the tree and estimate genetic distances with default settings. Tree robustness was assessed with 100 bootstrap replicates, and bootstrap values are shown in black adjacent to each node. Branch lengths are indicated in blue below each branch.

**Figure 3.**
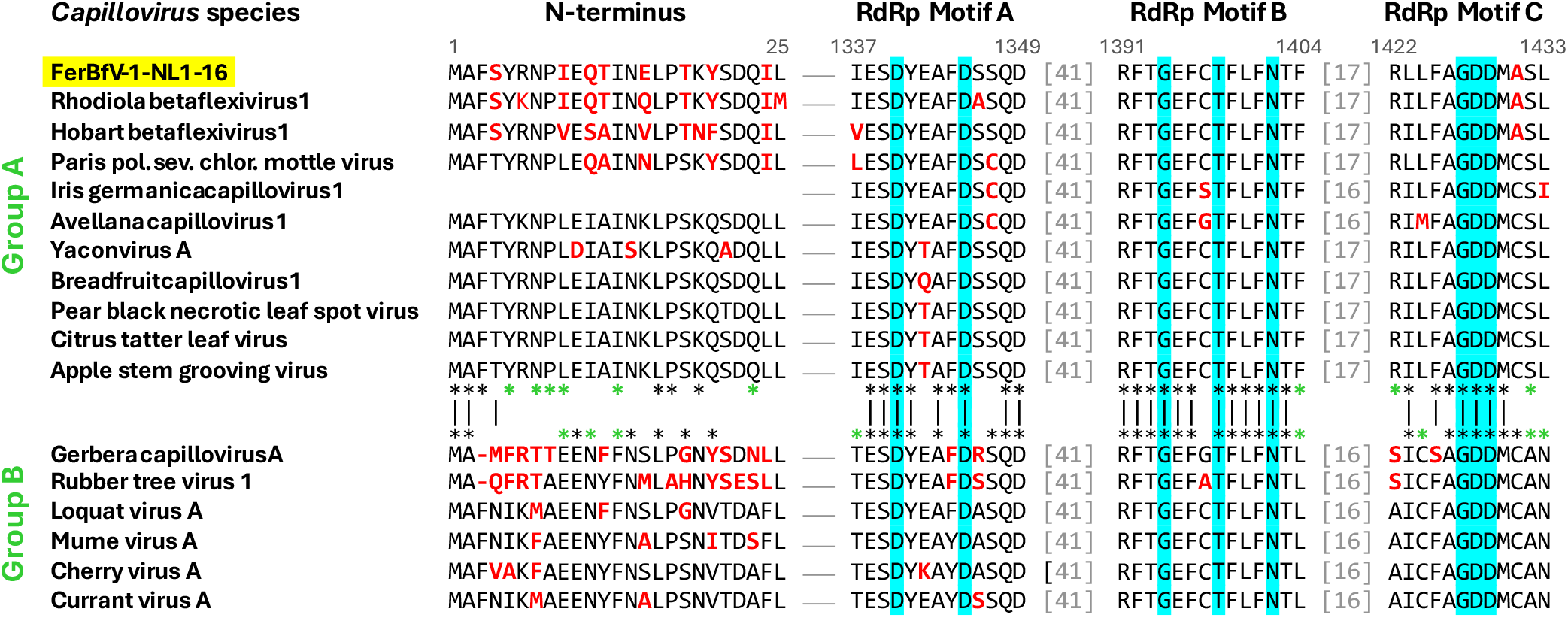
Comparison of conserved sequences in the *Capillovirus* ORF-1 polyprotein. Protein sequence alignment of the N-terminus and RdRp domain sequence motifs (A, B, and C) from representative members of the genus *Capillovirus* are shown. Amino acids that diverge from the consensus sequence for each *Capillovirus* group are highlighted in red. The top and bottom rows of asterisks represent conserved residues within groups A and B, respectively, with black vertical lines connecting conserved amino acids between both groups. Green asterisks denote residues that are distinctly conserved within one group and do not occur in the other group (Jia and Gong, 2019). Hallmark amino acids of the RdRp motifs are shaded blue.

## Concluding remarks

With the discovery of a new *Betaflexivirus* species, we added a tiny piece to the global virome puzzle. Although metagenomic analysis can generate new virus sequences at a much higher rate than our detailed approach, it often results in numerous partial virus sequences and frequently lacks detailed annotation. However, we feel that both approaches have their merits and that global virome research benefits from both.

Although the *Ferraria crispa* plants containing the FerBfV-1 virus exhibit a disease phenotype, it cannot be attributed to this virus as these plants also harbored several other viruses.

The fact that we easily detected several new virus species in the samples from a botanic garden (Dekker *et al*., 2024a and 2024b) supports our suggestion that these gardens may be rich reservoirs of plant viruses, many of them still unknown. This, plus the unique environment of (urban) botanic gardens, which could be considered ‘green islands’, have motivated us to conduct further follow-up experiments. These investigations promise not only more virus discoveries, but also valuable insights into virus evolution.

## Acknowledgements

We would like to express our sincere gratitude to Sarina Veldman, Martin Smit, Iris van Kleinwee and Reinout Havinga from the Hortus Botanicus in Amsterdam, The Netherlands, for their invaluable support in providing us with plant leaf material for this study. This research was directly and indirectly funded by the Swammerdam Institute for Life Sciences of the University of Amsterdam.

## Data availability

The raw sequence reads have been deposited in the NCBI Sequence Read Archive under BioProject accession number PRJNA1137160. The Ferraria betaflexivirus 1 isolate NL1-16 genome sequence has been deposited in NCBI GenBank under accession PQ073131.

## Supplemental information

**Supplemental Table ST1.** Read counts for mapping of smallRNA-seq and RNA-seq data to the Ferraria betaflexivirus-1 genomic RNA sequence. Read count indicates all reads that mapped to the Ferraria betaflexivirus-1 genome. Unique reads indicate the number of smallRNA-seq reads that mapped uniquely. Coverage indicates the breadth of coverage of the Ferraria betaflexivirus-1 genomic RNA sequence, expressed as a percentage. Coverages higher than 95% are indicated in yellow.

| Sample ID | Host plant genus | smallRNA-seq |  |  | RNAseq |
| --- | --- | --- | --- | --- | --- |
|  |  | Read count | Unique reads | Coverage (%) | Read count |
| S10 | *Albucca canadensis* | 219 | 112 | 1.9 | 624 |
| S23 | *Albucca canadensis* | 1,013 | 455 | 1.7 | 95 |
| S24 | *Albucca canadensis* | 1,209 | 515 | 1.7 | 499 |
| S05 | *Albucca canadensis* L FMLeight | 380 | 175 | 2.6 | 1,806 |
| S01 | *Drimiopsis maculata* Lindl Paxton | 1,851 | 527 | 1.8 | 1,281 |
| S14 | *Drimiopsis maculata* Lindl Paxton | 4,984 | 1,073 | 4.0 | 1,035 |
| S06 | *Ferraria crispa* Burm | 3,971 | 2,116 | 95.9 | 112,343 |
| S16 | *Ferraria crispa* Burm | 42,317 | 9,417 | 99.9 | 65,388 |
| S17 | *Ferraria crispa* Burm | 13,468 | 5,073 | 99.2 | 56,608 |
| S18 | *Ferraria crispa* Burm | 10,274 | 4,565 | 98.8 | 7,504 |
| S07 | *Ferraria crispa* Burm subsp *crispa* | 4,573 | 869 | 1.4 | 187 |
| S08 | *Ferraria crispa* Burm subsp *crispa* | 318 | 179 | 1.4 | 1,782 |
| S19 | *Ferraria crispa* Burm subsp *crispa* | 849 | 490 | 1.7 | 481 |
| S20 | *Ferraria crispa* Burm subsp *crispa* | 971 | 525 | 1.4 | 4,286 |
| S21 | *Ferraria crispa* Burm subsp *crispa* | 944 | 541 | 1.4 | 787 |
| S12 | *Gladiolus splendens* Sweet Herb | 250 | 190 | 1.6 | 802 |
| S15 | *Lachenalia aloides* Asch Graebert var *luteola* | 1,143 | 671 | 7.3 | 808 |
| S02 | *Lachenalia aloides* Asch Graebert var *quadricolor* | 246 | 77 | 1.4 | 1,127 |
| S03 | *Lachenalia bulbifera* Cirillo Engl | 97 | 73 | 1.6 | 1,148 |
| S04 | *Lachenalia bulbifera* Cirillo Engl | 55 | 47 | 1.3 | 1,160 |
| S25 | *Lachenalia corymbosa* JCManning Goldblatt | 180 | 134 | 1.4 | 1,048 |
| S26 | *Lachenalia kliprandensis* WF Barker | 258 | 176 | 1.4 | 220 |
| S09 | *Lachenalia liliiflora* Jacq | 551 | 262 | 1.3 | 595 |
| S22 | *Lachenalia stayneri* WFBarker | 475 | 289 | 1.7 | 346 |
| S11 | *Tritonia pallida* Ker Gawl subsp *pallida* | 47 | 42 | 1.2 | 213 |

**Supplemental Table ST2.**
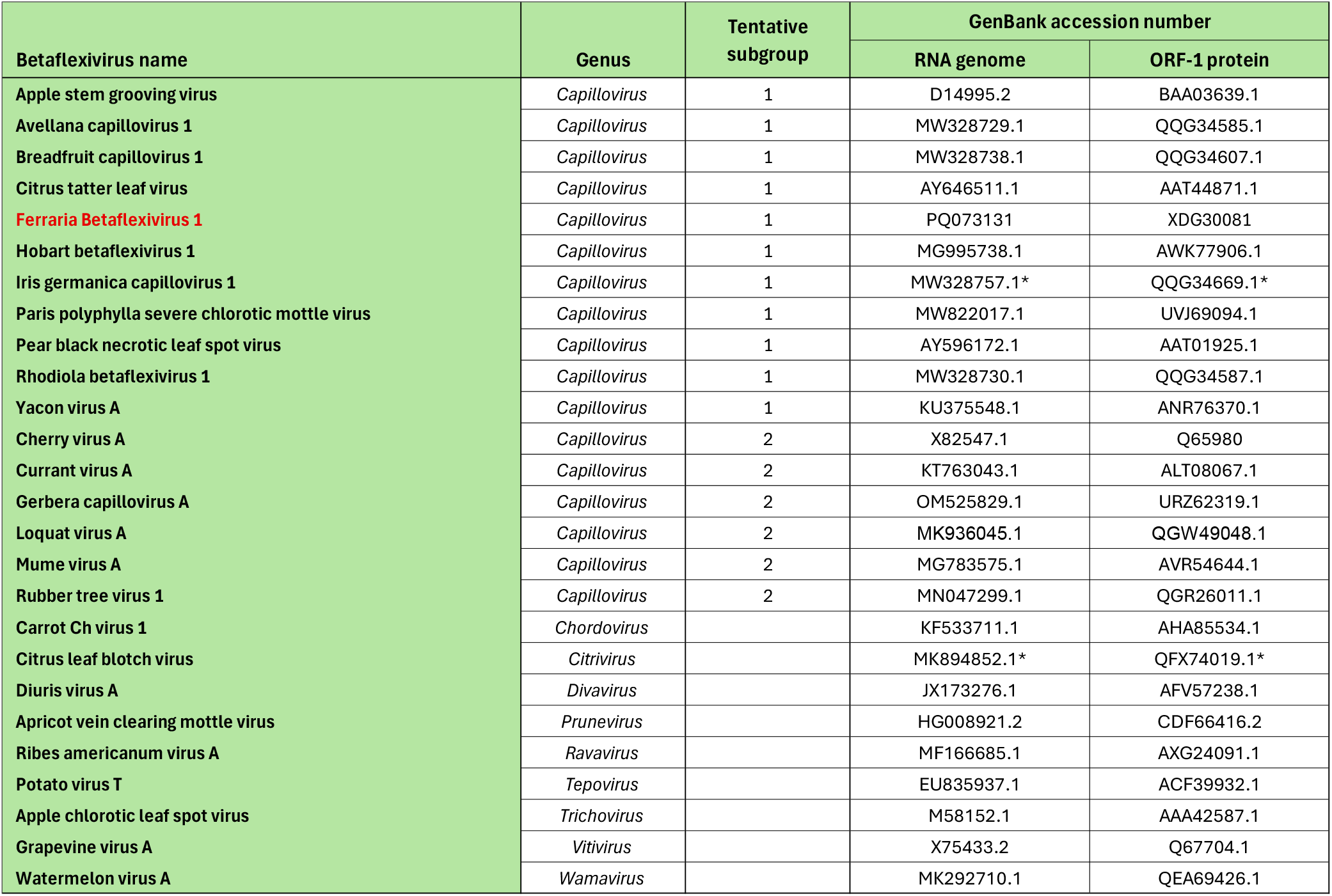
GenBank accession numbers for *Betaflexiviruses*. The new Ferraria Betaflexivirus 1 discovered in the current study is shown in red. Incomplete sequences are indicated with an asterisk.

| Betaflexivirus name | Genus | Tentative subgroup | GenBank accession number |  |
| --- | --- | --- | --- | --- |
|  |  |  | RNA genome | ORF-1 protein |
| Apple stem grooving virus | *Capillovirus* | 1 | D14995.2 | BAA03639.1 |
| Avellana capillovirus 1 | *Capillovirus* | 1 | MW328729.1 | QQG34585.1 |
| Breadfruit capillovirus 1 | *Capillovirus* | 1 | MW328738.1 | QQG34607.1 |
| Citrus tatter leaf virus | *Capillovirus* | 1 | AY646511.1 | AAT44871.1 |
| **Ferraria Betaflexivirus 1** | *Capillovirus* | 1 | PQ073131 | XDG30081 |
| Hobart betaflexivirus 1 | *Capillovirus* | 1 | MG995738.1 | AWK77906.1 |
| Iris germanica capillovirus 1 | *Capillovirus* | 1 | MW328757.1* | QQG34669.1* |
| Paris polyphylla severe chlorotic mottle virus | *Capillovirus* | 1 | MW822017.1 | UVJ69094.1 |
| Pear black necrotic leaf spot virus | *Capillovirus* | 1 | AY596172.1 | AAT01925.1 |
| Rhodiola betaflexivirus 1 | *Capillovirus* | 1 | MW328730.1 | QQG34587.1 |
| Yacon virus A | *Capillovirus* | 1 | KU375548.1 | ANR76370.1 |
| Cherry virus A | *Capillovirus* | 2 | X82547.1 | Q65980 |
| Currant virus A | *Capillovirus* | 2 | KT763043.1 | ALT08067.1 |
| Gerbera capillovirus A | *Capillovirus* | 2 | OM525829.1 | URZ62319.1 |
| Loquat virus A | *Capillovirus* | 2 | MK936045.1 | QGW49048.1 |
| Mume virus A | *Capillovirus* | 2 | MG783575.1 | AVR54644.1 |
| Rubber tree virus 1 | *Capillovirus* | 2 | MN047299.1 | QGR26011.1 |
| Carrot Ch virus 1 | *Chordovirus* |  | KF533711.1 | AHA85534.1 |
| Citrus leaf blotch virus | *Citrivirus* |  | MK894852.1* | QFX74019.1* |
| Diuris virus A | *Divavirus* |  | JX173276.1 | AFV57238.1 |
| Apricot vein clearing mottle virus | *Prunevirus* |  | HG008921.2 | CDF66416.2 |
| Ribes americanum virus A | *Ravavirus* |  | MF166685.1 | AXG24091.1 |
| Potato virus T | *Tepovirus* |  | EU835937.1 | ACF39932.1 |
| Apple chlorotic leaf spot virus | *Trichovirus* |  | M58152.1 | AAA42587.1 |
| Grapevine virus A | *Vitivirus* |  | X75433.2 | Q67704.1 |
| Watermelon virus A | *Wamavirus* |  | MK292710.1 | QEA69426.1 |

